# Machine learning identification of *Pseudomonas aeruginosa* strains from colony image data

**DOI:** 10.1101/2022.09.02.506375

**Authors:** Jennifer B. Rattray, Ryan J. Lowhorn, Ryan Walden, Pedro Márquez-Zacarías, Evgeniya Molotkova, Gabriel Perron, Claudia Solis-Lemus, Daniel Pimentel Alarcon, Sam P. Brown

## Abstract

When grown on agar surfaces, microbes can produce distinct multicellular spatial structures called colonies, which contain characteristic sizes, shapes, edges, textures, and degrees of opacity and color. For over one hundred years, researchers have used these morphology cues to classify bacteria and guide more targeted treatment of pathogens. Advances in genome sequencing technology have revolutionized our ability to classify bacterial isolates and while genomic methods are in the ascendancy, morphological characterization of bacterial species has made a resurgence due to increased computing capacities and widespread application of machine learning tools. In this paper, we revisit the topic of colony morphotype on the within-species scale and apply concepts from image processing, computer vision, and deep learning to a dataset of 69 environmental and clinical Pseudomonas aeruginosa strains. We find that colony morphology and complexity under common laboratory conditions is a robust, repeatable phenotype on the level of individual strains, and therefore forms a potential basis for strain classification. We then use a deep convolutional neural network approach with a combination of data augmentation and transfer learning to overcome the typical data starvation problem in biological applications of deep learning. Using a train/validation/test split, our results achieve an average validation accuracy of 92.9% and an average test accuracy of 90.7% for the classification of individual strains. These results indicate that bacterial strains have characteristic visual ‘fingerprints’ that can serve as the basis of classification on a sub-species level. Our work illustrates the potential of image-based classification of bacterial pathogens and highlights the potential to use similar approaches to predict medically relevant strain characteristics like antibiotic resistance and virulence from colony data.

**Author Summary:** Since the birth of microbiology, scientists have looked at the patterns of bacterial growth on agar (colony morphology) as a key tool for identifying bacterial species. We return to this traditional approach with modern tools of computer vision and deep learning and show that we can achieve high levels of classification accuracy on a within-species scale, despite what is considered a ‘data-starved’ dataset. Our results show that strains of the environmental generalist and opportunistic pathogen *Pseudomonas aeruginosa* have a characteristic morphological ‘fingerprint’ that enables accurate strain classification via a custom deep convolutional neural network. Our work points to extensions towards predicting phenotypes of interest (e.g. antibiotic resistance, virulence), and suggests that sample size limitations may be less restrictive than previously thought for deep learning applications in biology, given appropriate use of data augmentation and transfer-learning tools.

## Introduction

Since Semmelweis, Lister, and Koch began studying pure cultures of microorganisms, scientists have tried to organize the immense diversity of bacteria into orderly categories to understand their behavior, including pathogenesis. In particular, Koch pioneered the method of determining how specific microorganisms cause distinct diseases and highlighted the importance of laboratory culture for the identification and targeted treatment of pathogenic bacteria (1). By identifying the specific bacterial cause of disease, we can better manage and eradicate the infection. When a sample from an environment or clinical source is plated on solid media at sufficient dilution, individual colonies can be observed, where each colony grows from a single founding bacterial cell (Figure 1A). These clonal groups have characteristic sizes, shapes, edges, textures, and degrees of opacity and color (Figure 1B) (2). These characteristics describe the morphology of a single colony and formed the basis for the earliest attempts at categorization. In modern clinical laboratories, morphological tests play a role but are increasingly replaced by biochemical and genomic methods of classification (3).

**Figure 1.**
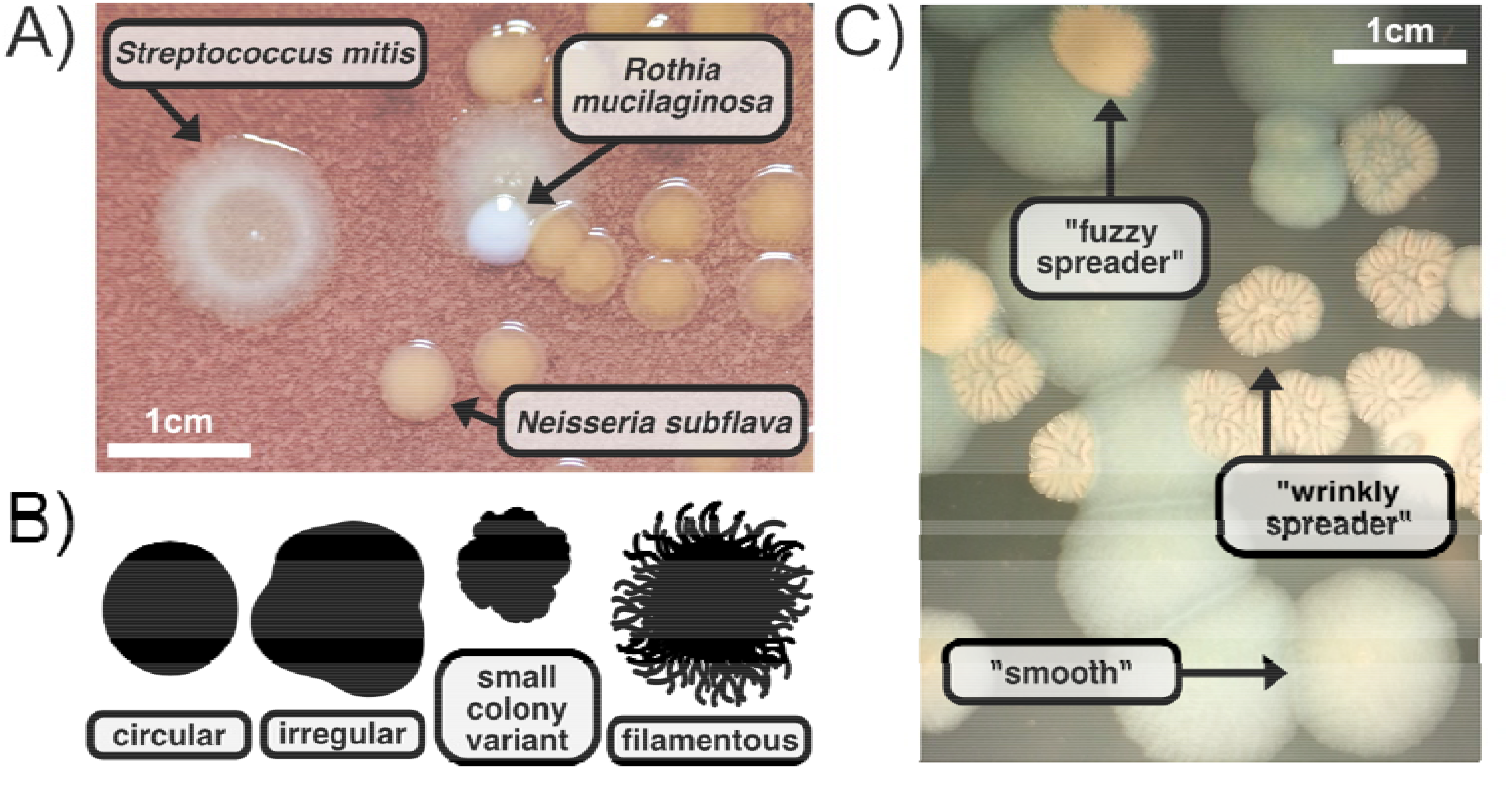
Bacterial colony morphology varies across and within species. A) Morphological identification of bacterial species from a mixed culture plated on Chocolate Agar: *Rothia mucilaginosa* (smallest white circles), *Neisseria subflava* (larger tan round circles), and *Streptococcus mitis* (large bullseye circles). B) A common example of 2D colony morphology features include the appearance of the colony edges. C) Morphological identification of bacterial strains, in this case called “smooth”, “wrinkly spreader”, and “fuzzy spreader”(4), from a culture containing only *Pseudomonas fluorescens* plated on King’s Medium B Agar.

While initial classification was morphology based, advances in genome sequencing technology have revolutionized our ability to classify bacterial isolates. Widespread use of 16S rRNA amplicon sequencing not only allows for robust taxon identification to the genus scale (5,6), but also fosters phylogenetic organization of bacterial taxa (7,8). Marker gene or whole genome sequencing provides finer scale strain identification and allows specific focus on markers of pathogenicity (via identification of virulence factors), drug resistance or other characteristics of interest. Transcriptomic sequencing provides a more direct window into the behavior of pathogens, to assay for example whether virulence factor or drug resistance genes are actually being expressed (9).

While genomic methods are in the ascendancy, morphological characterization of bacterial species has made a resurgence. How a population of bacteria grow is a direct result of the interplay between their genotype, phenotype, and environment (10) and *a priori* contains critical phenotypic information on the behavior of individual microbes. For example, colony morphotype can indicate specific mutations in a bacteria, e.g. small colony variants and cyclic di-GMP regulation (11) or rugosity and phenazine production (12). Morphological characterization and data science have collided to provide insights into image-based prediction of microbial taxa (13–18). The earliest attempts introduced light-scattering for presence-absence detection of microorganisms on surfaces (19,20), which paved the way for image-based identification of *Listeria* species from light scattering patterns of *Listeria* colonies (13). This developed into an automated light scattering sensor called BARDOT (<u>BA</u>cterial <u>R</u>apid <u>D</u>etection using <u>O</u>ptical scattering <u>T</u>echnology) that produced scatter patterns sensitive enough to distinguish between *Escherichia coli* serotypes (14), virulence mutants (15), and industrial *Staphylococcus* contaminants (16). At the same time, imaged-based prediction diversified to use more accessible standard light microscopes instead of specialized equipment, putting the focus on new analysis methods like colony morphology ontology (18) and deep learning (17,21).

Deep learning algorithms have gained popularity (17,21,22) as machine learning becomes ubiquitous across fields. Several supervised deep learning models have demonstrated near human accuracy in classification of various image datasets (23) and in the case of bacterial biofilms, deep learning has outperformed human characterization of single and mixed species biofilms (24). Zielinksi et al. (17) investigate several models for colony classification utilizing a deep convolutional neural network (D-CNN) on a rich dataset pre-trained on the ImageNet (25) dataset, achieving a very high validation accuracy of ∼ 97%. However, existing work using a D-CNN approach has focused on classification across species, not across strains within a species. Additionally, a ubiquitous challenge in many biology applications (including the one in this paper) is the limited number of samples. For example, the hallmark ImageNet image-based dataset (26–28) contains more than 14 million images across 1000 classes. Outside a colloquial ‘rule of thumb’ that you need at least 1000 samples per class, there is no absolute way to determine a ‘minimum’ sample size (29).

We focus on within species classification of *Pseudomonas aeruginosa*, a gram-negative bacterium capable of causing severe chronic illness and a major cause of acute hospital-acquired infections (30–33). While it is widely viewed as a generalist and can be isolated from a wide number of environments including water and soil, it is frequently found in human/animal-impacted environments (30). *P. aeruginosa* has large variety of mechanisms that allow it to respond to its environment (34–36). There are frequently differences in behavior across strains, like type and production of secreted products (37,38), which can result in observable morphological diversity when cultured on agar surfaces (39–43). Classic examples of morphological variation in P. *aeruginosa* caused by underlying genetic differences include small colony variants related to cyclic di-GMP regulation (11), rugosity related to phenazine production (12), and mucoidy related to secreted polymer production (44).

In this paper, we use a collection of 69 clinical and environmental *P. aeruginosa* isolates which present a range of phenotypic features (e.g. exopolysaccharide production, virulence gene expression, and antibiotic resistance profiles) that have been previously described in (45–47). From these 69 isolates we generate a library of 266 *P. aeruginosa* colony images in a training / validation dataset, plus a separately generated 69-image test dataset. As expected, we document a high level of variation in the physical appearance of colonies across strains. Additionally, we see that strain morphology in *P. aeruginosa* is consistent across replicates. We then use a D-CNN approach and a combination of data augmentation (48,49) and transfer learning (50) to classify strains of *P. aeruginosa* and achieve an average validation accuracy of 92.9% and an average test accuracy of 90.7%.

## Results

### Morphological variation across strains and replicates

Individual isolates from our 69-strain collection were spotted onto Luria-Bertani (LB) agar plates supplemented with Congo Red, a dye commonly used in microbiology to stain the biofilm matrix and extracellular polysaccharides (Figure 2). This was done in quadruplicate and generated a training and validation library of 266 colonies after quality control (10 colonies were removed due to debris or writing obscuring part of the image). Colonies were grown for 72 hours, which was determined during the pilot experiment to be enough time to reveal major morphological differences while minimizing outward expansion. This resulted in an imaged library of image colonies that ranged in size from 12M (approximately 0.74cm x 0.74cm) to 183M pixels (approximately 2.8cm x 2.8cm).

**Figure 2.**
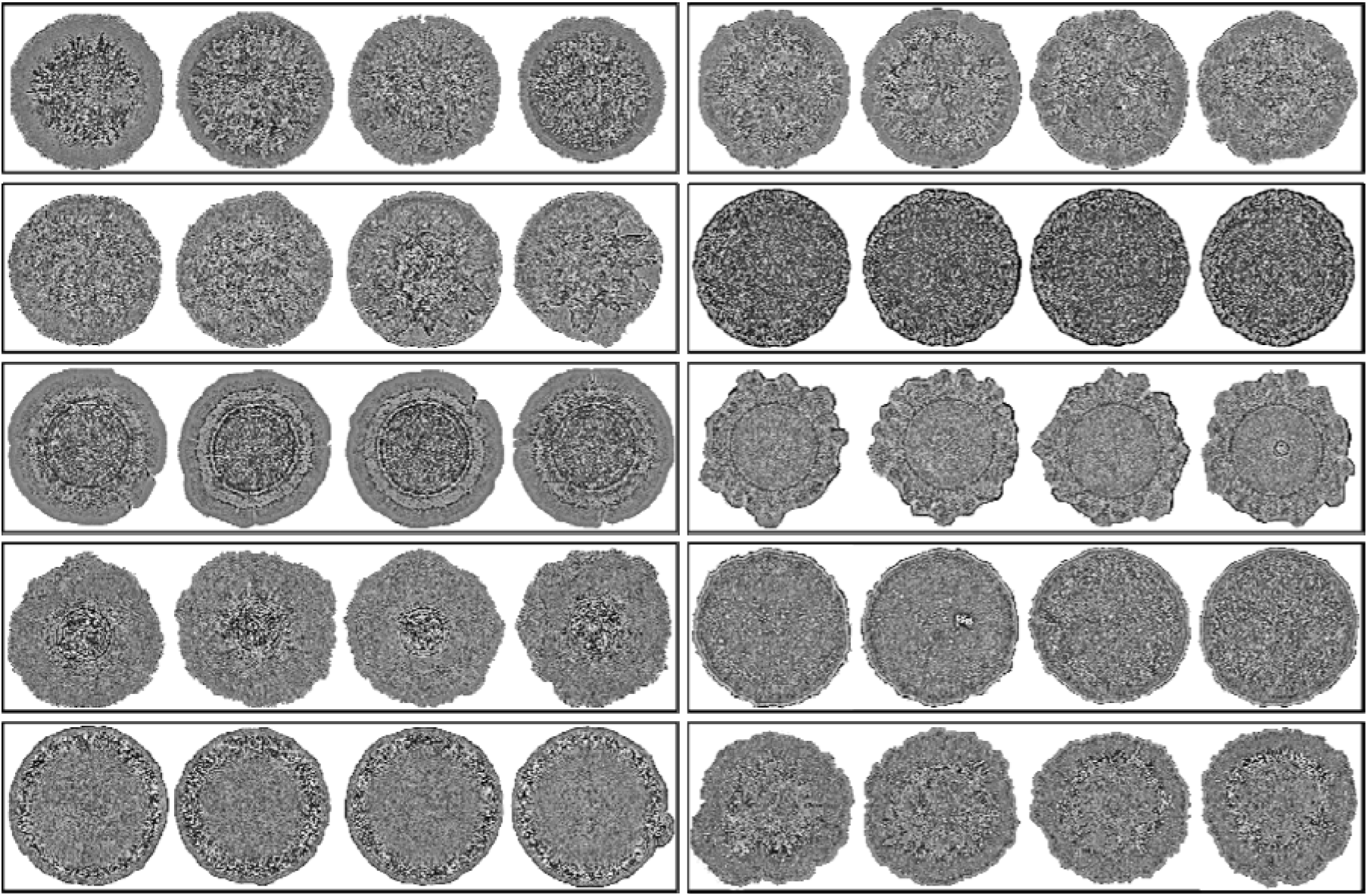
Random sample of the *P. aeruginosa* train/validation strain colony dataset. 10 strains (in four-fold replication) were selected randomly without replacement from a complete list of 69 strains via the native sample() command in R. From top to bottom: (left) strain 25, strain, 38, strain 52, strain 113, strain 1; (right) strain 127, strain 316, strain 298, strain 121, strain 174. Colony size has been adjusted for viewing purposes.

In our first approach to characterizing morphological diversity, we use 8 simple descriptive metrics (Figure 3A). Classic morphological descriptive variables include area, radius, perimeter, centroid (the point in which the three medians of the triangle imposed within the image intersect), eccentricity (the ratio of the distance between the foci of the ellipse and its major axis length), circularity (deviation of the perimeter from a superimposed perfect circle), bounding disk center (center of mass of the brightest pixels), and caliper diameter (the caliper dimension obtained by averaging over all orientations).

**Figure 3.**
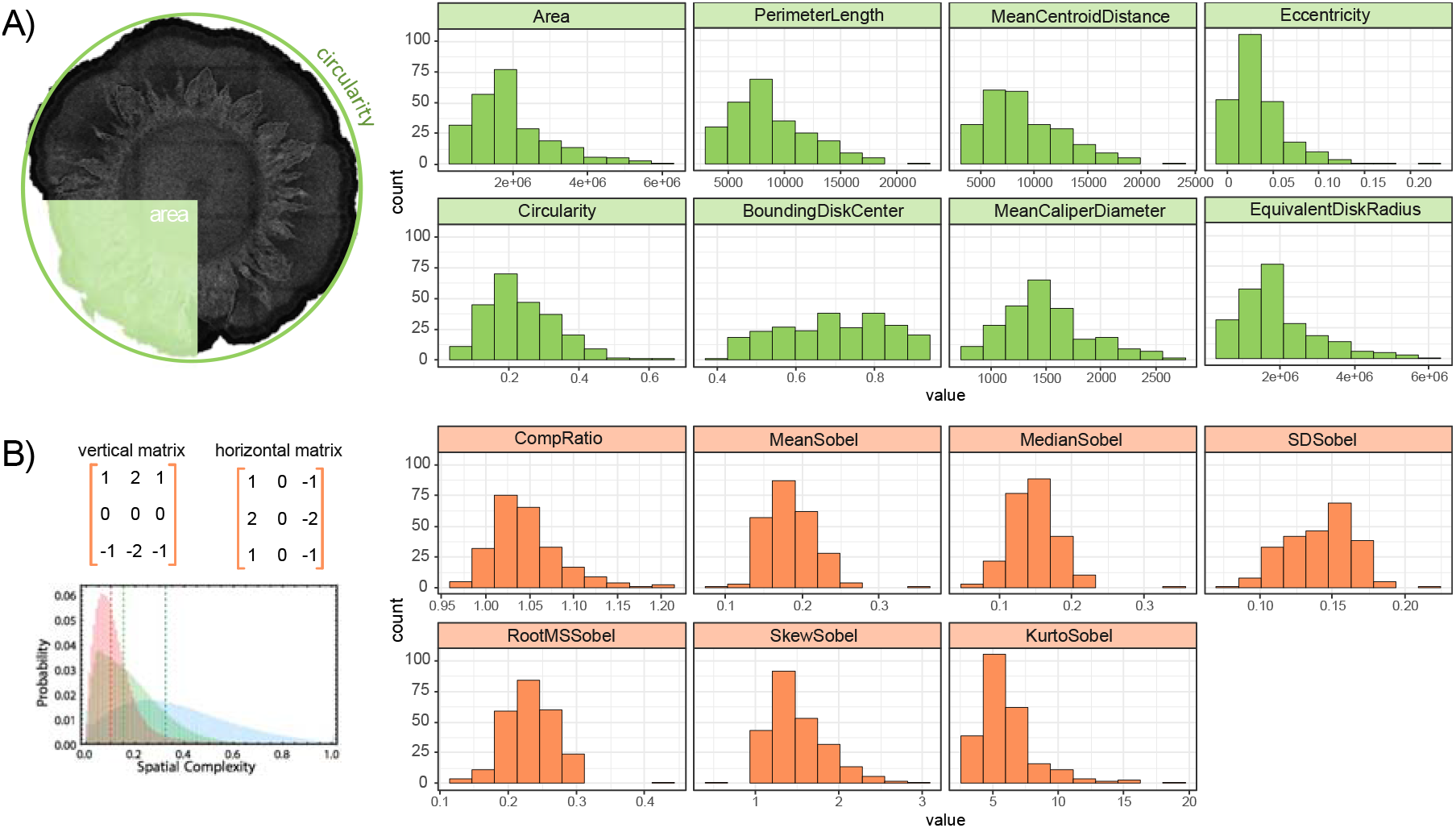
Diversity of *P. aeruginosa* strains in both classic morphological descriptive variables and derived complexity descriptive variables across 69 strains and 266 colonies. Histograms are built from all replicates of all strains. A) classic metrics used to describe colony appearance. B) Derived metrics from image processing and computer vision to describe colony complexity including compression ratio (relative reduction in size of data) and 6 descriptive statistics derived from the Sobel–Feldman operator.

To expand on these classic descriptive variables, we next introduce complexity metrics (Figure 3B) used in image processing and computer vision (51,52). Compression ratio is a single variable that describes how repetitive an image is, i.e. how repetitive are certain patterns in the pixel distribution and how many times do they occur? Specifically, it is a measurement of the reduction in size of pixel data produced by, in our case, the traditional JPEG compression algorithm. An alternate approach to quantifying image complexity is using the Sobel–Feldman operator, a popular method to detect edges in images due to its ability to detect gradients. At every pixel in an image, it calculates the pixel intensity of the vertical and horizontal adjacent pixels and expresses the results as vectors. You can then plot these gradient vectors creating a unique profile for every colony. These profiles can then be summarized by describing their distribution (mean, median, standard deviation, skew, root mean square, and kurtosis). Unsurprisingly, we find that many of these metrics are highly correlated (Supplemental Figure S1).

Figure 3 summarizes variation in our metrics across both strains and replicates. In Figure 4, we assess the extent of variation using coefficients of variation (CV = mean / standard deviation), across replicates (black bars) and across strains (grey bars). With the exception of the eccentricity and circularity metrics, we see coefficients of variation across replicates are low (standard deviation << mean), and less than the coefficient of variation across strains. This pattern of variation indicates that colony morphology under common laboratory conditions is a robust, repeatable phenotype on the level of individual strains, and therefore forms a potential basis for strain classification.

**Figure 4.**
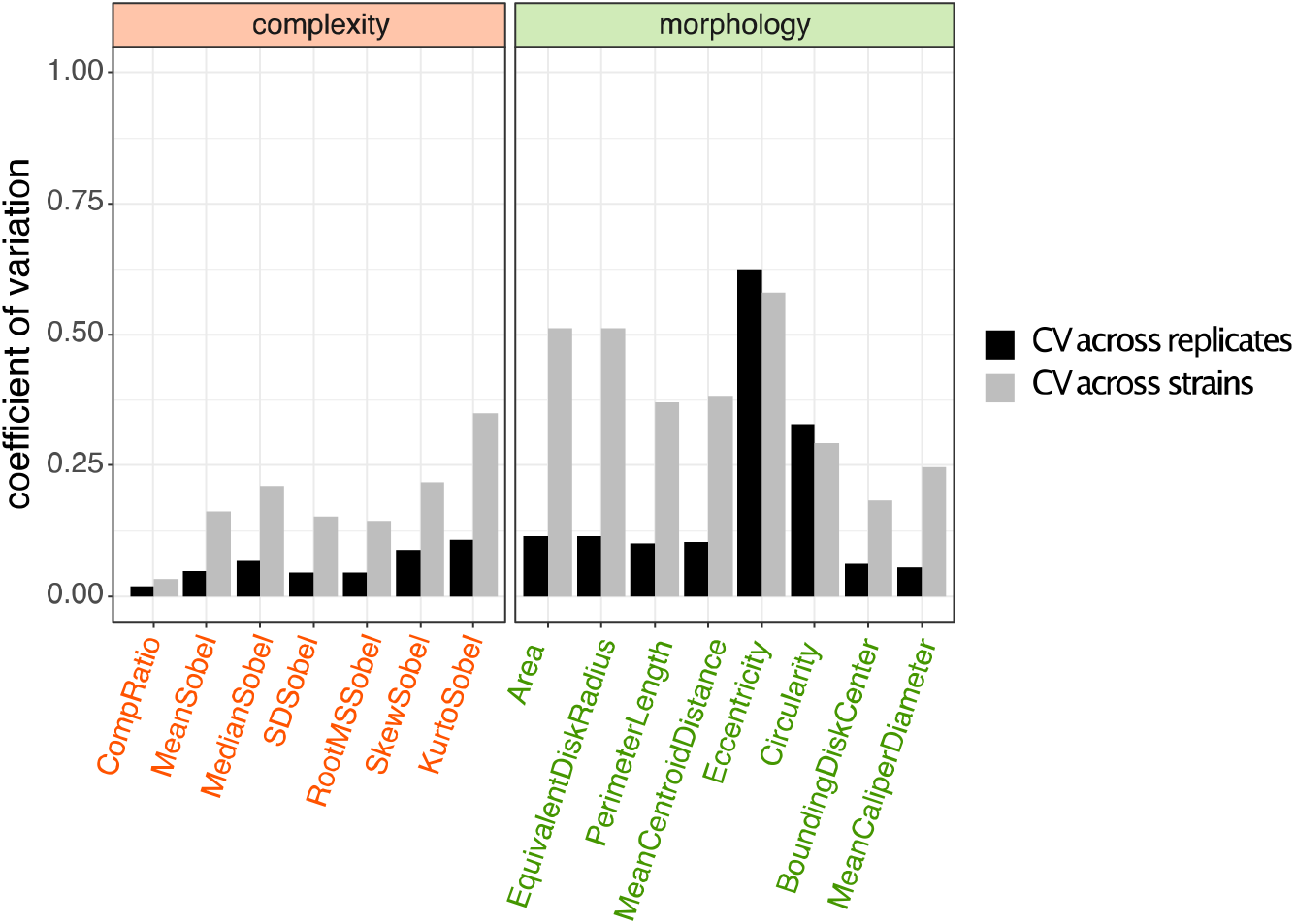
Morphological metrics are generally stable across replicates. Coefficient of variation (mean / standard deviation) across replicates (black) and across strains (grey). With the exception of the eccentricity and circularity metrics, coefficients of variation across replicates are low (standard deviation << mean), and less than the coefficient of variation across strains.

### Classifying strain identity from image data

Considering our initial results indicating robust morphological traits for each strain, we reasoned that morphological data could be used to classify colonies via a deep learning approach. Deep convolutional neural networks (D-CNNs) typically require larger datasets than those found in biology-usually a minimum of thousands of examples to train properly (53). To address this issue, we applied various transfer learning (50) and data augmentation techniques (48,49) (see methods for details). This initial dataset was split into 90% training and 10% validation sets.

High performance on our validation dataset provides some confidence that the CNN approach can successfully classify previously unseen strain images. Yet validation results merit caution, as the validation images were all generated on the same agar plate, imaged on the same day and from the same overnight culture. These shared features raise the risk that a CNN is detecting technical batch effects of the experimental environment (e.g. variations in the lighting conditions) rather than strain-specific features of interest. To address this concern we generated a separate 69-image test dataset (1 image per strain) using distinct overnight cultures and produced by a different experimenter on a different day (following the same protocol).

In a first round of computational experiments, we sought to compare the performance of four transfer learning models (Figure 5, Table S1), using our standard data preprocessing and augmentation choices (normalized pixel intensities; shear / zoom / flip / rotation / brightness / scaling data augmentation – see methods). To determine the best approach for our dataset, our performance metrics are accuracy (the number of correct predictions divided by the total number of predictions x100) and loss (a summation of the errors, calculated via cross-entropy, made for each sample). We chose to use cross-entropy for loss instead of negative log-likelihood since it automatically transforms the raw outputs of the neural network into probabilities (via a softmax activation).

**Figure 5:**
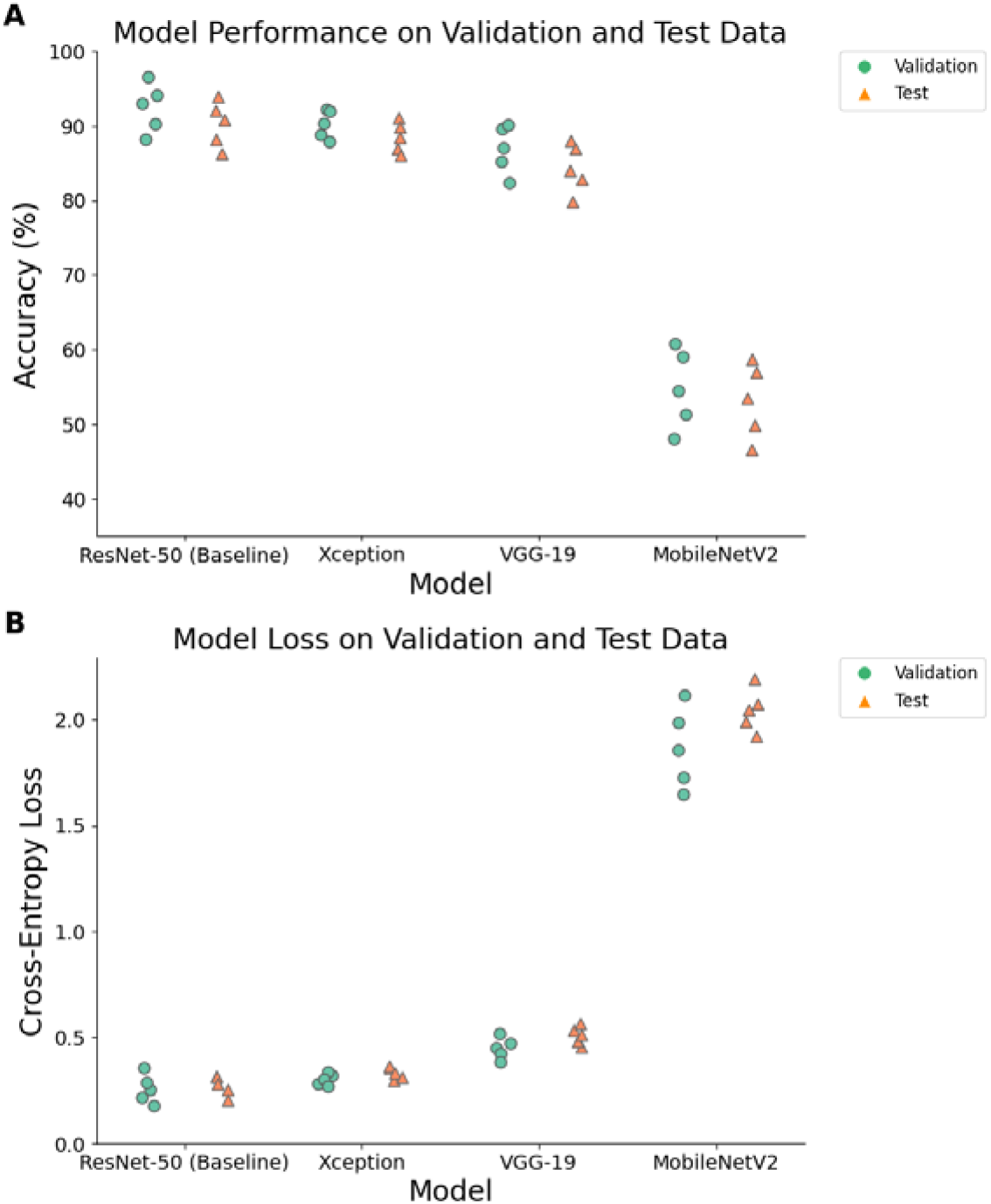
Performance comparison of transfer learning methods. The performance of four trained transfer learning models (ResNet-50, VGG-19, MobileNetV2 and Xception, see methods) were evaluated on both validation (blue circles) and test (orange triangles) datasets. Each computational experiment was replicated five-fold (five separate training runs for each model), allowing statistical comparison of approaches. (A) Accuracy scores (the number of correct predictions divided by the total number of predictions x100). (B) Loss scores (a summation of the errors made for each sample). Statistical comparisons are summarized in Table S1.

Figure 5 illustrates that model performance varies significantly between transfer learning models (e.g. for test accuracy: ANOVA, F = 124.2, df = 3, p < 0.05. See Table S1B).

MobileNetV2 performs significantly worse than the other transfer learning models on both accuracy and loss metrics (Tukey HSD, p < 0.05 on all comparisons, see Table S1C-F), while VGG-19 performs significantly worse than ResNet and Xception on test loss (Table S1D). ResNet and Xception do not show significantly different performance in posthoc tests (Table S1C-F). In terms of best individual model run performance, both the top accuracy and top loss was achieved by the ResNet model (Figure 5), so we use this as our baseline in our subsequent analyses.

In a second round of experiments, we take the best-performing model from Figure 5 (Resnet-50), and assess the contribution of data pre-processing, augmentation and training steps in generating the high levels of performance illustrated in Figure 5. Table 1 illustrates that removing each component of data augmentation leads to a significant drop in model accuracy (Tukey HSD, p < 0.05 on all comparisons, see Table S2B-E), and the loss of all augmentation leads to a substantial drop in performance (38.4 +/- 7.4% validation accuracy, 34.7 +/- 7.4% test accuracy). The removal of pre-trained weights results in a similar drop in performance.

**Table 1.** Performance contribution of data pre-processing, augmentation, and training. To assess contributions, we took the trained ResNet-50 model (Figure 5) as a baseline method and assessed the impact of removing components of our methods pipeline. Each computational experiment was replicated five-fold (five separate training runs for each model), allowing statistical comparison of approaches. Across replicates we report average (+/- standard deviation) accuracy (the number of correct predictions divided by the total number of predictions x100) and loss (a summation of the errors made for each sample). Statistical comparisons are summarized in Table S2.

| **Modification to baseline (ResNet)** | **Validation accuracy** | **Validation loss** | **Test accuracy** | **Test loss** |
| --- | --- | --- | --- | --- |
| Remove rotation augmentation | $76.83 \pm 7.061$ | $0.667 \pm 0.368$ | $72.12 \pm 7.061$ | $0.707 \pm 0.408$ |
| Remove brightness augmentation | $78.33 \pm 6.982$ | $0.618 \pm 0.346$ | $52.17 \pm 7.038$ | $1.622 \pm 0.456$ |
| Remove shift augmentation | $69.27 \pm 7.716$ | $0.789 \pm 0.429$ | $65.13 \pm 7.716$ | $0.829 \pm 0.469$ |
| Remove shear augmentation | $70.84 \pm 7.431$ | $0.761 \pm 0.416$ | $66.25 \pm 7.431$ | $0.801 \pm 0.456$ |
| Remove horizontal flip augmentation | $72.58 \pm 7.287$ | $0.735 \pm 0.403$ | $67.85 \pm 7.287$ | $0.775 \pm 0.443$ |
| Remove zoom augmentation | $71.92 \pm 7.415$ | $0.754 \pm 0.413$ | $67.08 \pm 7.415$ | $0.794 \pm 0.453$ |
| Remove all augmentation | $38.42 \pm 7.360$ | $2.302 \pm 0.468$ | $34.75 \pm 7.360$ | $2.372 \pm 0.508$ |
| No pre-trained weights | $32.81 \pm 6.822$ | $2.302 \pm 0.368$ | $29.08 \pm 7.234$ | $2.742 \pm 0.541$ |
| Remove image normalization | $73.52 \pm 7.534$ | $0.722 \pm 0.397$ | $68.39 \pm 7.534$ | $0.762 \pm 0.437$ |

The performance of our best performing ResNet-50 model (Figure 5) indicates that while most classification calls are accurate, there remain a number of mistaken calls in our validation tests. To look more closely at these errors, we present in Figure 6 a reduced confusion matrix showing all classification errors across our five replicate runs of the ResNet-50 model. Of the 69 strains considered in this study, only 5 were not classified with perfect precision. Strains 2 and 84 were the most challenging for our model, attaining the lowest test accuracies of 40% and 20% respectively. In contrast, 64 out 69 strains were classified with perfect accuracy across 5 independent model runs.

**Figure 6.**
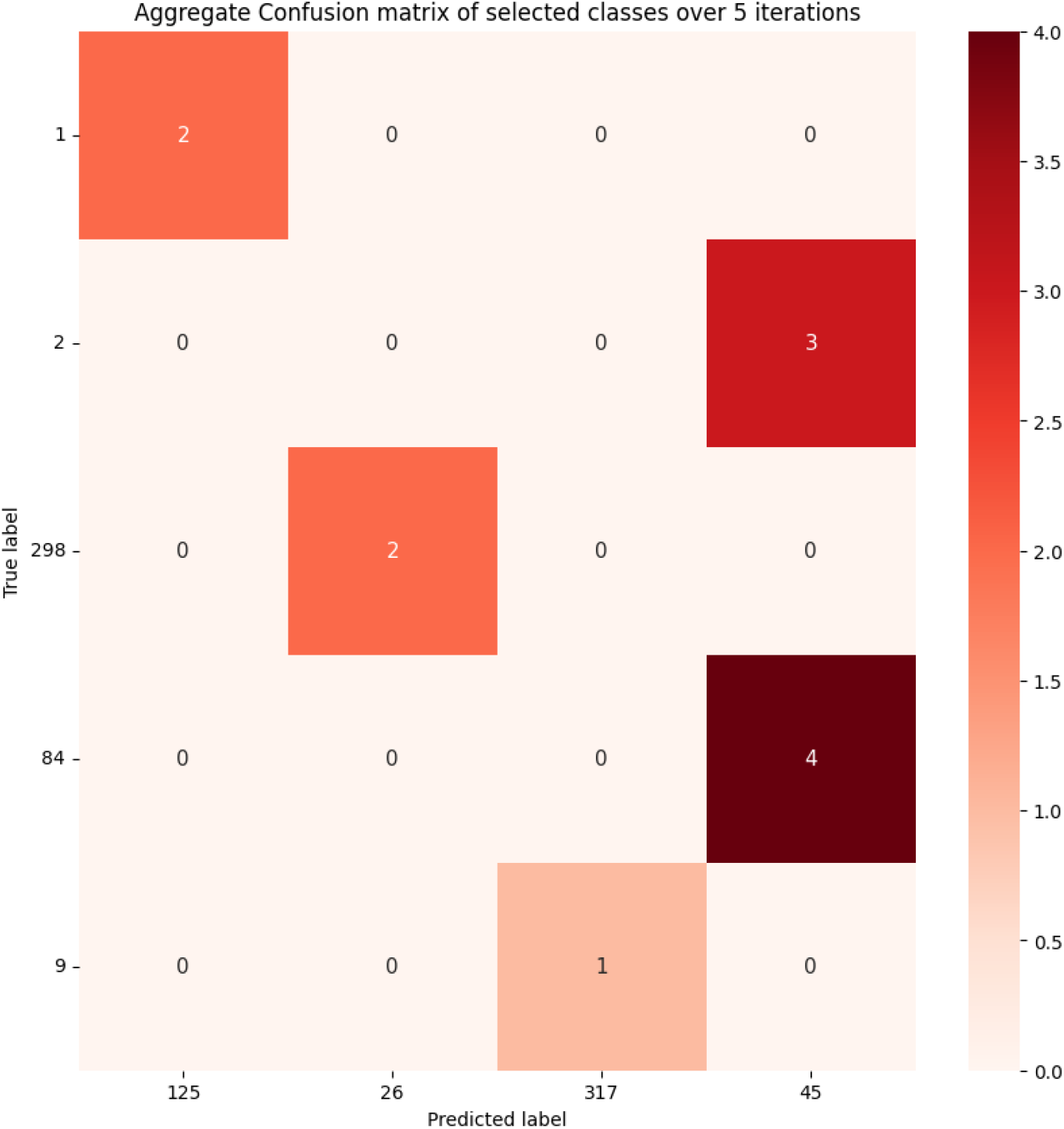
Reduced confusion matrix, aggregating all test data errors across five iterations of the ResNet-50 model. Of the 69 strains in this study, only 5 strains were not classified with 100% precision (the 5 rows). For brevity, this matrix includes only the strains that were misclassified by our model instead of the whole 69 x 69 confusion matrix.

Our results in Tables 1 and 2 show that D-CNN models are capable of effective bacterial colony classification down to the sub-species (strain) level, and that performance is dependent on details of data pre-processing and augmentation. This success does not rule out that other simpler models can also effectively classify strains. To establish simple model benchmarks, we next evaluate the performance of ‘shallow learning’ classification models (support vector machines, SVMs), trained on our colony metric features (Figures 3, 4), or on features extracted by our deep learning ResNet 50 model (Table 2). These benchmarking results highlight that SVM tools cannot match the performance of our D-CNN models (Figure 5), whether they are trained on *a priori* identified colony metric features (Figures 3,4) or on features extracted from our successful D-CNN models.

**Table 2.** Performance comparison with shallow learning (SVM) models. We contrast validation and test accuracy for our trained ResNet model (see also Figure 5, Table S1) with accuracy metrics for radial basis function (RBF) SVMs trained on colony metric data (Figure 3,4) and on features extracted from the trained ResNet model. See methods for details of SVM models and feature extraction. Statistical comparisons are summarized in Table S3.

| model | Validation accuracy (%) | Test accuracy (%) |
| --- | --- | --- |
| ResNet 50 | 92.93 ± 4.831 | 90.73 ± 5.073 |
| Colony metric data, RBF SVM | 46.73 ± 7.532 | 33.43 ± 6.341 |
| ResNet 50 feature extraction and RBF SVM | 65.29 ± 1.754 | 62.75 ± 1.931 |

## Discussion

Our results show that *P. aeruginosa* strains have a characteristic morphological ‘fingerprint’ (Figures 2-4) that enables accurate strain classification via a custom deep convolutional neural network (Figure 5). When trained on morphological data, our most successful model can classify strain identity with a test accuracy that exceeds 90% despite what is considered a “data starved” dataset. Our work points to extensions towards predicting phenotypes of interest (e.g. antibiotic resistance, virulence), and suggests that sample size limitations may be less restrictive than previously thought for deep learning applications in biology. In the following paragraphs we will review these implications, while addressing caveats and next steps for this research.

The existence of a morphological ‘fingerprint’ that can be classified via machine learning architecture opens the potential to relate strain identity to key phenotypes of interest, such as antimicrobial resistance (AMR) or virulence. While species differ in their levels and types of virulence (54), there are also significant differences in virulence between strains of the same species (55,56). In so far as AMR or virulence is a stable and repeatable property of a strain, our classification model could simply “look up” the recorded values in a dataset based on the strain prediction. Yet this “look up” approach is vulnerable to variation within strains due for instance to gain or loss in virulence factor and AMR genes, or mutations in the regulatory control of these genes.

To address the challenge of predicting phenotypes such as virulence or AMR, a future direction would be to extend our deep learning approach to directly predict phenotypes of interest, either through a classification (qualitative trait) or regression (quantitative trait) framework. Our current analysis shows that morphological ‘fingerprints’ are sufficient to identify individual strains, but this does not directly imply that we can use image data to successfully predict disease traits (analogously, human fingerprints are sufficient to identify individuals, but are not considered predictive of human disease traits). We speculate that colony ‘fingerprints’ are likely to allow for trait prediction, due to paths of common causality from bacterial modes of growth to colony morphology and disease phenotypes. Given the established connection between biofilm growth and antibiotic tolerance (57), it is possible that our default growth settings (Figure 2) will reveal differences in patterning that correlate with differences in antibiotic resistance (and/or tolerance). We further anticipate that our approach would become more discriminatory if imaging data was also collected in the presence/absence of relevant stressors of interest (e.g. antibiotics or immune effectors).

Deep learning methods are commonly described as ‘black box’ methodologies (58), presenting challenges for model interrogation and mathematical justification for sample size requirements. Even when classification through deep learning has a monotonically increasing prediction accuracy, the nature of the image collection process in biological research will limit the total size of the dataset. This is a contrast to machine learning applications in other fields that have continually and often exponentially increasing datasets (e.g., spending habits and surveillance cameras). Considering these challenges, arguably our best alternatives for increasing the size of biological datasets for image-based prediction in microbiology are data augmentation and transfer learning, which we combined in this study. Our algorithm transfers learning from the canonical ImageNet dataset while also using standard data augmentation techniques (rotations, shifts, zooms, shears, flips, reflections, brightness.). Future tests to explore the amount of data needed for accurate classification could be run by reducing the size of the dataset by decreasing the amount of data augmentation performed or by reducing the number of replicate colonies used. Moving forward, generative adversarial networks, a class of machine learning frameworks that learns to generate new data by competing two neural networks, is promising technique that could supplement current data augmentation efforts (59,60).

Prior to our data augmentation, we downsampled the original dataset in order to match the image dimensions of the ImageNet dataset which we pretrained on and to minimize computational time. The original dataset contains colony images of 12 million to 183 million pixels (depending on colony size) while our downsized colony images each contained a standardized 60 thousand pixels. The success of strain classification based on these heavily down-sampled images suggests that classification is possible based only on more ‘macro’ scale morphological features that are still discriminable at this resolution. Yet this down-sampling is essentially throwing away a rich array of more micro-scale image features (Figure 7), and so opens the possibility for future analyses to explore these features. The inset panels in Figure 7 illustrate striking local and finer-scale features that are present across strains. These panels suggest an additional data augmentation option, where each colony image is processed into multiple 60K chunks, sampling at different locations within the image and at different spatial scales. Comparing model performance using 60K images taken at different spatial scales could provide valuable information on what are the most diagnostic spatial scales.

**Figure 7.**
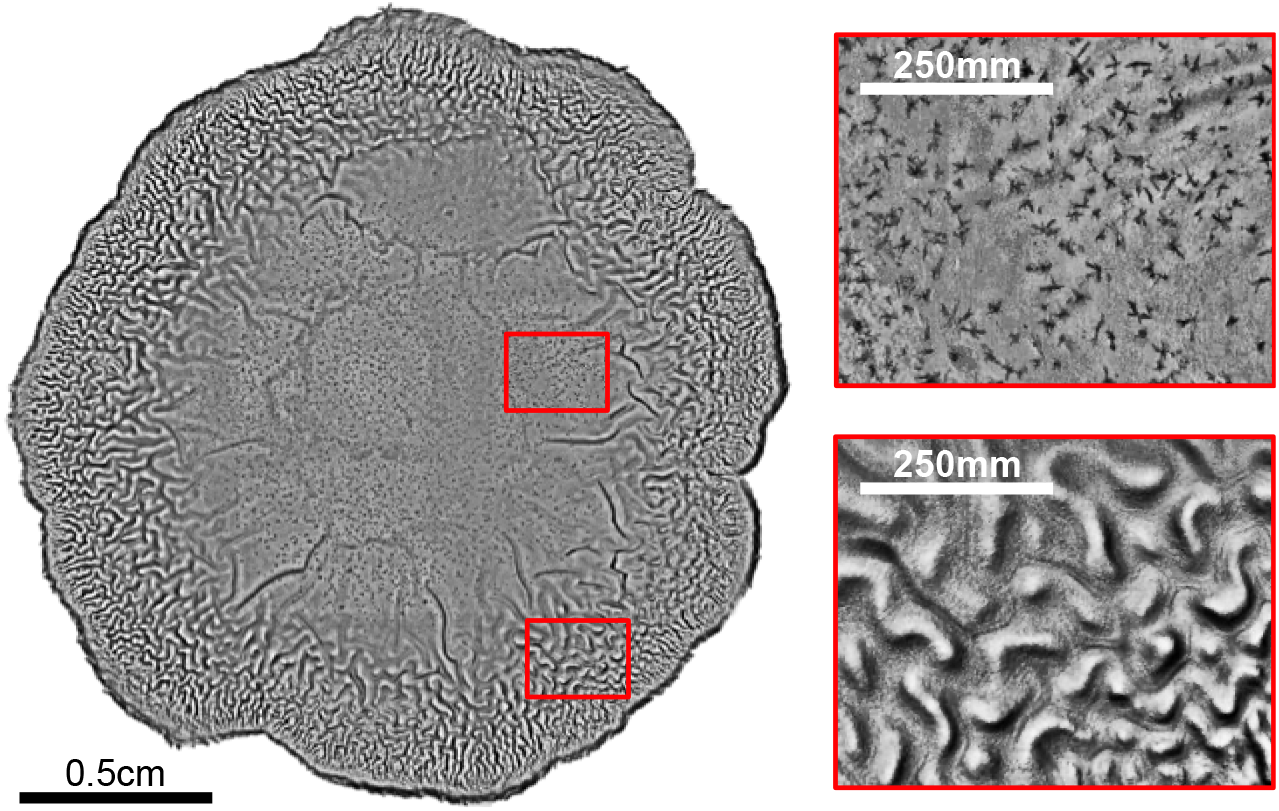
Avenues for finer scale image segmentation and augmentation. Insets reveal finer-scale features at smaller spatial scales.

Our analysis to date implicitly assumes that our image data represent independent samples of *P. aeruginosa* morphology, which due to phylogenetic relationships among these strains is not the case. A simple null expectation given these phylogenetic relationships is that strains that are most morphologically similar (i.e., similar morphological metrics, Figure 3; and more commonly confused in classification, Figure 6) will tend to have a shorter phylogenetic distance. Under this phylogenetic null model, strains with minimal phylogenetic differences will have overlapping morphological metrics and will not provide dependable classification results. We do not expect this pattern to hold uniformly, as we know that single-gene changes can generate large-scale changes to colony morphology (11,12,44). Future work can systematically explore the impacts of single gene mutations through automated screening of transposon mutagenesis libraries (61), potentially aiding in the discovery of previously unrecognized genes involved in colony / biofilm formation. We also encourage future work applying our general methodology to sets of strains in other species, as we expect our approach will generalize beyond the model organism *P. aeruginosa*. In summary, the level of classification accuracy achieved in this work illustrates the potential of image-based prediction tools, deep learning, transfer learning, and data augmentation in the characterization of bacterial strains.

## Methods

### Strain Culturing

This study uses a collection of 69 clinical and environmental *P. aeruginosa* isolates (45–47), each grown and imaged in quadruplicate. We used Luria-Bertani (LB) agar (1.5%) plates supplemented with 0.08% Congo Red, a dye commonly used to characterize biofilm formation which binds to compounds in the extracellular biofilm matrix. In an effort to minimize variation in plate preparation and the age of the plates, 20ml volume plates were made 24 hours before each experiment with a large, motorized pipette and kept in a sealed plastic sleeve at 4C. Strains were inoculated into LB broth and incubated shaking overnight at 37C. The day of the experiment, each plate was then sliced into 4 sections using a sterilized razor and tweezers, which both limited colony outgrowth and prevented the diffusion of small molecules between colonies. 5µl of the overnight culture was then spotted onto pre-prepared, sectioned LB agar Congo Red plates. After the spots dried, plates were parafilmed to retain moisture and placed in a 37C static incubator. All strains were incubated at 37C for 72 hours, which was determined during the pilot experiment to be enough time to reveal major morphological differences while minimizing outward expansion.

### Colony Imaging

After 72 hours, strains were imaged on a Nikon Eclipse Ti inverted microscope at 4X using the DS-Fi2 colored camera. This results in a pixel-to-size ratio is 2.07uM per pixel. Large, stitched images were required to scan an entire colony and were generated with Nikon’s built-in software. Some replicate colonies had to be excluded from analysis due to error (strains 13, 47, 78, 86, 122, 124, 132, 210 have 3 replicates and strain 329 has 2 replicates). This resulted in a dataset of 266 *P. aeruginosa* colony images that ranged in size from 12,644,652 pixels (approximately 0.74cm x 0.74cm) to 183,329,280 pixels (approximately 2.8cm x 2.8cm).

### Morphological and Complexity Metrics

For the morphological and complexity measurements, colony images were analyzed in Mathematica. Images were scaled down by a factor of four, to make computations timelier, and the backgrounds were cropped out by using a watershed transformation, binarization, and selection of the contiguous non-zero area surrounding the colony. The morphological metrics (Area, EquivalentDiskRadius, PerimeterLength, MeanCentroidDistance, Eccentricity, Circularity, BoundingDiskCenter, and MeanCaliperDiameter) were all calculated using the native Mathematica commands. For the complexity metrics, CompRatio was calculated by comparing the number of bytes in the uncompressed and compressed (using the native Mathematica command) colony images. A Sobel-Freidman operator was applied in both x and y dimensions and Sobel histograms were generated. MeanSobel, MedianSobel, SDSobel, RootMSSobel, SkewSobel, and KurtosisSobel were all then calculated using the native Mathematica commands.

### Data pre-processing for machine learning

Our experimental pipeline is an integration of several steps: preprocessing, transfer learning, and data augmentation techniques, all aimed at maximizing the predictive performance of our deep learning model. In the preprocessing phase, all images are first downsampled to a uniform size of 224×224 pixels, consistent with the image dimensions used in the pretrained ImageNet database. To preserve the original aspect ratio and minimize distortions during the resizing process, we employed inter-area interpolation.

In addition to resizing, another important step in our preprocessing is image normalization. This includes two primary aspects. The first is the correction of image backgrounds: the algorithm iterates over the images and replaces any black pixels with white ones. This process aids in standardizing the image backgrounds across our dataset, ensuring that the model does not get biased by variations in the background and thus enhancing its robustness to different imaging conditions.

The second aspect of image normalization involves rescaling the pixel intensities. Raw images are composed of pixels with intensity values that usually fall within a range of 0 to 255. These large values may slow down the learning process and make the model more susceptible to suboptimal solutions. To mitigate this, we applied a transformation to rescale the pixel values to a 0-1 range. This process, often termed pixel normalization, can help improve the efficiency of the learning process by preventing large input values from causing numerical instability and by ensuring that the input features have a similar data distribution, thus assisting the optimization algorithm to converge more quickly (62).

Overall, these preprocessing steps serve as an integral part of the pipeline, shaping the data into a more suitable form for the model to learn effectively and efficiently, while ensuring robustness to potential sources of variation.

### Data augmentation

In order to further optimize our model’s performance and ensure its robustness, we utilized various data augmentation techniques. This approach, known to improve the generalization capability of deep learning models by artificially expanding the training dataset (63), involves making stochastic transformations to the training images, thereby mimicking variations likely to be seen in the real-world application of the model. One such transformation was brightness augmentation, where the brightness of the image is varied randomly within a range. Specifically, the brightness was altered by factors between 0.2 and 0.8, effectively simulating various lighting conditions that the images might be subjected to in real-world scenarios (64). This rescaling process not only aids in normalizing the data but also helps in mitigating the risk of gradient explosion, a phenomenon that can hinder the learning process (65). Random rotations within a range of -20 to 20 degrees and random shifts in image width and height by up to 20% of the original size are also used. These techniques simulate changes in the object’s orientation and position within the frame, thus allowing our model to learn invariance to such alterations (66). The application of shear transformations and zoom operations, both by up to 20%, mimic the effects of perspective distortion and changes in the object-camera distance respectively. These techniques enhance the model’s ability to recognize the subject regardless of variations in the camera perspective or object scale (49). Finally, we also incorporated horizontal flipping of images, a transformation especially useful for our dataset since it does not possess any inherent left-right bias. This method doubles the size of the dataset and helps the model generalize better by learning symmetrical features (67).

### Train/validation/test split

Images from the initial 4 replicate imaging experiment were divided (following augmentation) into a 90-10 train/validation split. The imaging protocol was repeated by a separate experimenter to generate an independent test sample.

### Transfer learning models

In our study, we employed four deep learning models to classify bacterial strains, leveraging the advantages of both transfer learning and architectural diversity. Each model used was pre-trained on the ImageNet dataset and further fine-tuned to our specific task. This section details the deep learning models used, the modifications we made, and our rationale for these choices.

Before the training of any models, the topmost layer of each pre-trained model was removed, and replaced by a new flatten layer. This was followed by two pairs of dense and dropout layers, and finally, a softmax dense output layer. The dense layers were designed using Rectified Linear Unit (ReLU) activations (68), while the final dense layer employs a softmax activation function. This transforms the output into a probability distribution across 69 neurons, each corresponding to a unique strain identification within our dataset. The resulting confidence vector is obtained when the output layer is activated.

The decision to downsize the new dense layers from the original size (e.g., from 2048 to 1024 in ResNet-50) was influenced by the size of our dataset in contrast to ImageNet. Dropout layers were also incorporated to prevent overfitting, following the transfer learning research conducted by (69).

#### ResNet-50

ResNet-50 (70) is known for its residual blocks, which mitigate the vanishing and exploding gradient issues common in deep learning models. In our application, the weights of the pre-existing layers were frozen, and only the newly added dense layers were trained - a method known as ‘feature extraction.’ This method exploits the pre-trained model’s ability to extract generalized features, while the added layers learn to make predictions based on these features.

The use of ResNet-50 in bacterial strain identification tasks is well-documented. A study (71) employed a ResNet-based approach to classify three species of gram-positive bacteria, achieving an accuracy of 81%. Another study (72) compared the performance of various deep learning models, including ResNet, for bacilli detection, demonstrating the efficacy of these architectures in bacterial identification tasks. These previous successful uses of ResNet-50 provide a robust motivation for its selection in our study.

#### MobileNetV2

We employed the MobileNetV2 model for its compactness and efficiency, making it suitable for mobile and embedded vision applications (73). Similar to ResNet-50, we used the feature extraction approach with this model.

#### VGG-19

The VGG-19 model (53) was also utilized in our study due to its proven efficacy in various classification tasks. We implemented the feature extraction strategy, leveraging the learned features from the model and fine-tuning our custom layers for our specific task.

#### Xception

The Xception model was incorporated due to its unique architecture, designed to recognize nuanced patterns (74). For Xception, we adopted a ‘fine-tuning’ strategy, training all layers of the network with a smaller learning rate. This allowed the model to adjust the learned weights more accurately to our problem.

#### Benchmark Models

For comparison, two ‘shallow learning’ methods were used as benchmarks. First, a Radial Basis Function (RBF) Support Vector Machine (SVM) was employed for strain classification using colony metric data (Figure 3), following the successful application demonstrated by Chen et al. (75) in bacterial classification.

Additionally, a hybrid approach was adopted to exploit the strengths of both deep and shallow learning. Here, we extracted feature vectors from our top-performing deep learning model, ResNet-50, which were then used as input for a RBF SVM. This process of feature extraction involves using ResNet-50’s pre-trained weights to generate a representative vector of the input image. These vectors essentially encapsulate the critical visual patterns recognized by the model, thus serving as robust predictors in the SVM. This hybrid approach involves loading the pre-trained ResNet-50 model with weights trained on the ImageNet dataset and setting the ‘include_top’ parameter to False. This configuration returns a model that does not include the final dense layer, making it suitable for feature extraction. An image is loaded and preprocessed to match the input requirements of ResNet-50, and the model’s ‘predict’ function is then used to generate the feature vector. This feature vector is then used as input to the SVM for classification. This transfer learning strategy has been successfully applied in numerous studies, including Rahmayuna et al. (76), who achieved a remarkable classification accuracy of 90.33%.

## Acknowledgements

We thank Jennifer Farrell for culturing the mixed community in Figure 1 and members of the Center for Microbial Dynamics and Infection (CMDI), in particular Dr. Peter Yunker, Dr. Conan Zhao, and Dr. Yifei Wang, for valuable comments and discussion on this work. This research was supported by the National Science Foundation Graduate Research Fellowship Program under grant no. DGE-1650044, the Foundation for the National Institutes of Health grant no. 1R21AI143296, and the Foundation for the National Institutes of Health grant no. 1R21AI156817.

## Supporting Information

**Supplemental Figure S1.**
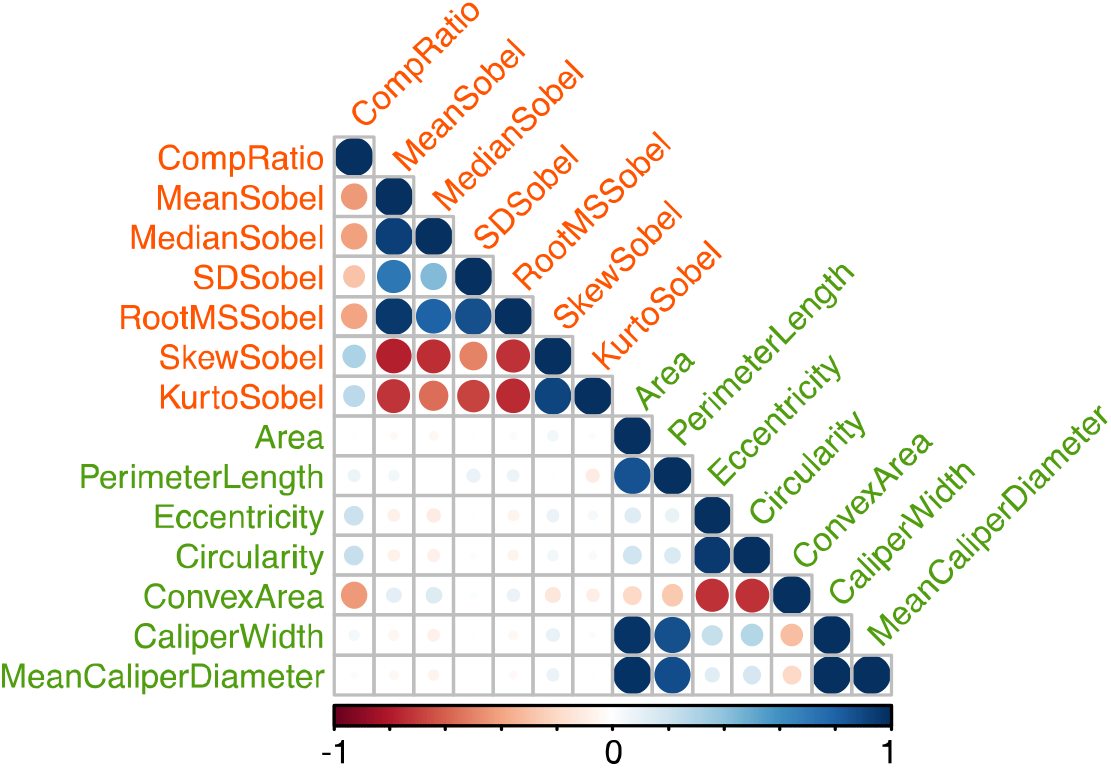
Correlation matrix for all morphological and complexity metrics. Morphological (green) and complexity (orange) metrics.

**Supplemental Table S1. Statistical comparisons of transfer learning methods (Figure 5).** The performance of four trained transfer learning models (ResNet-50, VGG-19, MobileNetV2 and Xception, see methods) were evaluated on both validation and test datasets. Each computational experiment was replicated five-fold (five separate training runs for each model), allowing statistical comparison of approaches.

**Table S1A.** Summary statistics. Across replicates we report average (+/- standard deviation) accuracy (the **number of correct predictions divided by the total number of predictions x100) and loss (**a summation of the errors made for each sample).

| Transfer learning Method | Validation accuracy (%) | Validation loss | Test accuracy (%) | Test loss |
| --- | --- | --- | --- | --- |
| ResNet 50 | 92.93 ± 4.831 | 0.251 ± 0.174 | 90.73 ± 5.073 | 0.278 ± 0.183 |
| VGG-19 | 86.96 ± 4.033 | 0.448 ± 0.130 | 83.92 ± 4.232 | 0.510 ± 0.137 |
| MobileNetV2 | 54.40 ± 6.252 | 1.853 ± 0.303 | 53.39 ± 6.563 | 2.043 ± 0.318 |
| Xception | 90.25 ± 4.561 | 0.300 ± 0.201 | 88.36 ± 4.788 | 0.330 ± 0.210 |

**Table S1B.** ANOVA table.

| ANOVA |  | df | F | P-value |
| --- | --- | --- | --- | --- |
| Validation Accuracy | 4700.077255 | 3.0 | 119.531973 | 3.627413e-11 |
| Test Accuracy | 4571.131895 | 3.0 | 124.245285 | 2.698834e-11 |
| Validation Loss | 8.867851 | 3.0 | 269.70656 | 6.616010e-14 |
| Test Loss | 10.635474 | 3.0 | 942.817267 | 3.337569e-18 |

**Tables S1C-F.** Post-hoc pairwise tests (Tukey HSD with alpha = 0.05).

| Supplementary Table S1C: Post Hoc Analysis for Validation Loss |  |  |  |  |  |  |
| --- | --- | --- | --- | --- | --- | --- |
| group1 | group2 | meandiff | p-adj | lower | upper | reject |
| MobileNetV2 | ResNet-50 (Baseline) | -1.6076 | 0.0 | -1.797 | -1.4182 | True |
| MobileNetV2 | VGG-19 | -1.4158 | 0.0 | -1.6052 | -1.2264 | True |
| MobileNetV2 | Xception | -1.5636 | 0.0 | -1.753 | -1.3742 | True |
| ResNet-50 (Baseline) | VGG-19 | 0.1918 | 0.0467 | 0.0024 | 0.3812 | True |
| ResNet-50 (Baseline) | Xception | 0.044 | 0.9088 | -0.1454 | 0.2334 | False |
| VGG-19 | Xception | -0.1478 | 0.1567 | -0.3372 | 0.0416 | False |
| **Supplementary Table 1D: Post Hoc Analysis for Test Loss** |  |  |  |  |  |  |
| **group1** | **group2** | **meandiff** | **p-adj** | **lower** | **upper** | **reject** |
| MobileNetV2 | ResNet-50 (Baseline) | -1.7704 | 0.0 | -1.8814 | -1.6594 | True |
| MobileNetV2 | VGG-19 | -1.5342 | 0.0 | -1.6452 | -1.4232 | True |
| MobileNetV2 | Xception | -1.7116 | 0.0 | -1.8226 | -1.6006 | True |
| ResNet-50 (Baseline) | VGG-19 | 0.2362 | 0.0001 | 0.1252 | 0.3472 | True |
| ResNet-50 (Baseline) | Xception | 0.0588 | 0.4513 | -0.0522 | 0.1698 | False |
| VGG-19 | Xception | -0.1774 | 0.0016 | -0.2884 | -0.0664 | True |
| **Supplementary Table 1E: Post Hoc Analysis for Validation Accuracy** |  |  |  |  |  |  |
| **group1** | **group2** | **meandiff** | **p-adj** | **lower** | **upper** | **reject** |
| MobileNetV2 | ResNet-50 (Baseline) | 37.688 | 0 | 31.1371 | 44.2389 | True |
| MobileNetV2 | VGG-19 | 32.128 | 0 | 25.5771 | 38.6789 | True |
| MobileNetV2 | Xception | 35.502 | 0 | 28.9511 | 42.0529 | True |
| ResNet-50 (Baseline) | VGG-19 | -5.56 | 0.1115 | -12.1109 | 0.9909 | False |
| ResNet-50 (Baseline) | Xception | -2.186 | 0.7763 | -8.7369 | 4.3649 | False |
| VGG-19 | Xception | 3.374 | 0.4751 | -3.1769 | 9.9249 | False |
| **Supplementary Table 1F: Post Hoc Analysis for Test Accuracy** |  |  |  |  |  |  |
| **group1** | **group2** | **meandiff** | **p-adj** | **lower** | **upper** | **reject** |
| MobileNetV2 | ResNet-50 (Baseline) | 37.13 | 0 | 30.7933 | 43.4667 | True |
| MobileNetV2 | VGG-19 | 31.206 | 0 | 24.8693 | 37.5427 | True |
| MobileNetV2 | Xception | 35.342 | 0 | 29.0053 | 41.6787 | True |
| ResNet-50 (Baseline) | VGG-19 | -5.924 | 0.0711 | -12.2607 | 0.4127 | False |
| ResNet-50 (Baseline) | Xception | -1.788 | 0.8501 | -8.1247 | 4.5487 | False |
| VGG-19 | Xception | 4.136 | 0.2801 | -2.2007 | 10.4727 | False |

**Supplemental Table S2. Performance contribution of data pre-processing, augmentation and training.** To assess contributions we took the trained ResNet-50 model (Figure 5, Table S1A) as a baseline method, and assessed the impact of removing components of our methods pipeline. Five-fold replicated results are summarized in Table 1.

**Supplemental Table S2A.** ANOVA table of data in table 1.

| ANOVA | sum_sq | df | F | P-value |
| --- | --- | --- | --- | --- |
| Validation Accuracy | 14795.48945 | 9.0 | 278.010813 | 3.655742e-33 |
| Test Accuracy | 14972.050258 | 3.0 | 241.435997 | 5.808255e-32 |
| Validation Loss | 30.322865 | 3.0 | 782.2722 | 4.790182e-42 |
| Test Loss | 40.431932 | 3.0 | 977.413531 | 5.719771e-44 |

**Supplemental Table S2B-E.** Post-hoc pairwise tests (Tukey HSD with alpha = 0.05).

| Table S2B. Post Hoc Analysis for Validation Loss |  |  |  |  |  |  |
| --- | --- | --- | --- | --- | --- | --- |
| Group 1 | Group 2 | Meandiff | p-adj | Lower | Upper | Reject |
| No pre-trained weights | Resnet-50 | -2.0578 | 0 | -2.1968 | -1.9188 | TRUE |
| Remove all augmentation | Resnet-50 | -0.457 | 0 | -0.596 | -0.318 | TRUE |
| Remove brightness augmentation | Resnet-50 | -0.3572 | 0 | -0.4962 | -0.2182 | TRUE |
| Remove horizontal flip augmentation | Resnet-50 | -2.0254 | 0 | -2.1644 | -1.8864 | TRUE |
| Remove image normalization | Resnet-50 | -0.457 | 0 | -0.596 | -0.318 | TRUE |
| Remove rotation augmentation | Resnet-50 | -0.3844 | 0 | -0.5234 | -0.2454 | TRUE |
| Remove shear augmentation | Resnet-50 | -0.4578 | 0 | -0.5968 | -0.3188 | TRUE |
| Remove shift augmentation | Resnet-50 | -0.5098 | 0 | -0.6488 | -0.3708 | TRUE |
| Remove zoom augmentation | Resnet-50 | -2.0628 | 0 | -2.2018 | -1.9238 | TRUE |

| Group 1 | Group 2 | Meandiff | p-adj | Lower | Upper | Reject |
| --- | --- | --- | --- | --- | --- | --- |
| No pre-trained weights | Resnet-50 | -2.5482 | 0 | -2.6917 | -2.4047 | TRUE |
| Remove all augmentation | Resnet-50 | -0.532 | 0 | -0.6755 | -0.3885 | TRUE |
| Remove brightness augmentation | Resnet-50 | -1.3998 | 0 | -1.5433 | -1.2563 | TRUE |
| Remove horizontal flip augmentation | Resnet-50 | -2.082 | 0 | -2.2255 | -1.9385 | TRUE |
| Remove image normalization | Resnet-50 | -0.532 | 0 | -0.6755 | -0.3885 | TRUE |
| Remove rotation augmentation | Resnet-50 | -0.4308 | 0 | -0.5743 | -0.2873 | TRUE |
| Remove shear augmentation | Resnet-50 | -0.5532 | 0 | -0.6967 | -0.4097 | TRUE |
| Remove shift augmentation | Resnet-50 | -0.609 | 0 | -0.7525 | -0.4655 | TRUE |
| Remove zoom augmentation | Resnet-50 | -2.5482 | 0 | -2.6917 | -2.4047 | TRUE |

| Group 1 | Group 2 | Meandiff | p-adj | Lower | Upper | Reject |
| --- | --- | --- | --- | --- | --- | --- |
| No pre-trained weights | Resnet-50 | 59.004 | 0 | 53.8552 | 64.1528 | TRUE |
| Remove all augmentation | Resnet-50 | 52.004 | 0 | 46.8552 | 57.1528 | TRUE |
| Remove brightness augmentation | Resnet-50 | 12.416 | 0 | 7.2672 | 17.5648 | TRUE |
| Remove horizontal flip augmentation | Resnet-50 | 17.94 | 0 | 12.7912 | 23.0888 | TRUE |
| Remove image normalization | Resnet-50 | 16.888 | 0 | 11.7392 | 22.0368 | TRUE |
| Remove rotation augmentation | Resnet-50 | 13.704 | 0 | 8.5552 | 18.8528 | TRUE |
| Remove shear augmentation | Resnet-50 | 19.738 | 0 | 14.5892 | 24.8868 | TRUE |
| Remove shift augmentation | Resnet-50 | 21.358 | 0 | 16.2092 | 26.5068 | TRUE |
| Remove zoom augmentation | Resnet-50 | 18.738 | 0 | 13.5892 | 23.8868 | TRUE |

| Group 1 | Group 2 | Meandiff | p-adj | Lower | Upper | Reject |
| --- | --- | --- | --- | --- | --- | --- |
| No pre-trained weights | Resnet-50 | 62.322 | 0 | 56.7641 | 67.8799 | TRUE |
| Remove all augmentation | Resnet-50 | 55.274 | 0 | 49.7161 | 60.8319 | TRUE |
| Remove brightness augmentation | Resnet-50 | 38.774 | 0 | 33.2161 | 44.3319 | TRUE |
| Remove horizontal flip augmentation | Resnet-50 | 22.868 | 0 | 17.3101 | 28.4259 | TRUE |
| Remove image normalization | Resnet-50 | 21.726 | 0 | 16.1681 | 27.2839 | TRUE |
| Remove rotation augmentation | Resnet-50 | 18.506 | 0 | 12.9481 | 24.0639 | TRUE |
| Remove shear augmentation | Resnet-50 | 24.6 | 0 | 19.0421 | 30.1579 | TRUE |
| Remove shift augmentation | Resnet-50 | 26.584 | 0 | 21.0261 | 32.1419 | TRUE |
| Remove zoom augmentation | Resnet-50 | 23.264 | 0 | 17.7061 | 28.8219 | TRUE |

**Supplemental Table S3. Performance comparison with shallow learning (SVM) models.** We contrast validation and test accuracy for our trained ResNet model (see also Table 1) with accuracy metrics for radial basis function (RBF) SVMs trained on colony metric data (Figure 3,4) and on features extracted from the trained ResNet model. See methods for details of SVM models and feature extraction.

**Supplemental Table S3A:** ANOVA of Table 2.

| ANOVA | sum_sq | df | F | P-value |
| --- | --- | --- | --- | --- |
| Validation Accuracy | 5268.1030 | 9.0 | 324.059336 | 3.608738e-11 |
| Test Accuracy | 7869.885293 | 3.0 | 909.536437 | 7.922344e-14 |

**Supplemental Table S3B-E.** Post-hoc pairwise tests (Tukey HSD with alpha = 0.05).

|  |
| --- |
| Table S3B. Post Hoc Analysis for Validation Accuracy: |

| Group 1 | Group 2 | Mean Difference | p-adj | Lower | Upper | Reject |
| --- | --- | --- | --- | --- | --- | --- |
| Colony metric data, RBF SVM | ResNet-50 | 45.73 | 0 | 40.9195 | 50.5405 | TRUE |
| Colony metric data, RBF SVM | ResNet 50 feature extraction and RBF SVM | 19.4 | 0 | 14.5895 | 24.2105 | TRUE |
| ResNet 50 feature extraction and RBF SVM | ResNet-50 | 26.33 | 0 | 21.5195 | 31.1405 | TRUE |

| Group 1 | Group 2 | Mean Difference | p-adj | Lower | Upper | Reject |
| --- | --- | --- | --- | --- | --- | --- |
| Colony metric data, RBF SVM | ResNet-50 | 56.094 | 0 | 52.5844 | 59.6036 | TRUE |
| Colony metric data, RBF SVM | ResNet 50 feature extraction and RBF SVM | 29.078 | 0 | 25.5684 | 32.5876 | TRUE |
| ResNet 50 feature extraction and RBF SVM | ResNet-50 | 27.016 | 0 | 23.5064 | 30.52 | TRUE |

## References

1. Koch R. Ueber den augenblicklichen Stand der bakteriologischen Choleradiagnose. Zeitschrift für Hygiene und Infektionskrankheiten 1983 14:1 [Internet]. 1893 Dec [cited 2022 Jul 19];14(1):319–38. Available from: https://link.springer.com/article/10.1007/BF02284324

2. Madigan MT, Brock TD. Brock biology of microorganisms. 12th editi. Biology of Microorganisms. San Francisco, CA: Pearson/Benjamin Cummings; 2009.

3. Bergey D, Krieg NR, Holt JG. Bergey’s manual of systematic bacteriology. Baltimore, MD: Williams & Wilkins; 1984.

4. Rainey PB, Travisano M. Adaptive radiation in a heterogeneous environment. Nature 1998 394:6688 [Internet]. 1998 Jul 1 [cited 2022 Jul 18];394(6688):69–72. Available from: https://www.nature.com/articles/27900

5. Woo PCY, Lau SKP, Teng JLL, Tse H, Yuen KY. Then and now: use of 16S rDNA gene sequencing for bacterial identification and discovery of novel bacteria in clinical microbiology laboratories. Clin Microbiol Infect [Internet]. 2008 [cited 2022 Jul 19];14(10):908–34. Available from: https://pubmed.ncbi.nlm.nih.gov/18828852/

6. Yarza P, Yilmaz P, Pruesse E, Glöckner FO, Ludwig W, Schleifer KH, et al. Uniting the classification of cultured and uncultured bacteria and archaea using 16S rRNA gene sequences. Nat Rev Microbiol [Internet]. 2014 [cited 2022 Jul 19];12(9):635–45. Available from: https://pubmed.ncbi.nlm.nih.gov/25118885/

7. Woese CR. Bacterial evolution. Microbiol Rev [Internet]. 1987 Jun [cited 2022 Jul 19];51(2):221. Available from: https://www.ncbi.nlm.nih.gov/pmc/articles/PMC373105/

8. Woese C. The universal ancestor. Proceedings of the National Academy of Sciences [Internet]. 1998 Jun 9 [cited 2022 Jul 19];95(12):6854–9. Available from: https://www.pnas.org/doi/abs/10.1073/pnas.95.12.6854

9. Güell M, Yus E, Lluch-Senar M, Serrano L. Bacterial transcriptomics: what is beyond the RNA horiz-ome? Nature Reviews Microbiology 2011 9:9 [Internet]. 2011 Aug 12 [cited 2022 Jul 19];9(9):658–69. Available from: https://www.nature.com/articles/nrmicro2620

10. Kalziqi A, Yanni D, Thomas J, Ng SL, Vivek S, Hammer BK, et al. Immotile Active Matter: Activity from Death and Reproduction. Phys Rev Lett [Internet]. 2018 Jan 5 [cited 2022 Jul 17];120(1):018101. Available from: https://journals.aps.org/prl/abstract/10.1103/PhysRevLett.120.018101

11. Starkey M, Hickman JH, Ma L, Zhang N, de Long S, Hinz A, et al. Pseudomonas aeruginosa rugose small-colony variants have adaptations that likely promote persistence in the cystic fibrosis lung. J Bacteriol [Internet]. 2009 Jun [cited 2016 May 18];191(11):3492–503. Available from: http://www.ncbi.nlm.nih.gov/pubmed/19329647

12. Recinos DA, Sekedat MD, Hernandez A, Cohen TS, Sakhtah H, Prince AS, et al. Redundant phenazine operons in Pseudomonas aeruginosa exhibit environment-dependent expression and differential roles in pathogenicity. Proc Natl Acad Sci U S A [Internet]. 2012 Nov 20 [cited 2022 Jul 3];109(47):19420–5. Available from: www.pnas.org/cgi/doi/10.1073/pnas.1213901109

13. Banada PP, Guo S, Bayraktar B, Bae E, Rajwa B, Robinson JP, et al. Optical forward-scattering for detection of Listeria monocytogenes and other Listeria species. Biosens Bioelectron. 2007 Mar 15;22(8):1664–71.

14. Tang Y, Kim H, Singh AK, Aroonnual A, Bae E, Rajwa B, et al. Light Scattering Sensor for Direct Identification of Colonies of Escherichia coli Serogroups O26, O45, O103, O111, O121, O145 and O157. PLoS One [Internet]. 2014 Aug 19 [cited 2022 Jul 17];9(8). Available from: /pmc/articles/PMC4138183/

15. Singh AK, Sun X, Bai X, Kim H, Abdalhaseib MU, Bae E, et al. Label-free, non-invasive light scattering sensor for rapid screening of Bacillus colonies. J Microbiol Methods. 2015 Feb 1;109:56–66.

16. Alsulami TS, Zhu X, Abdelhaseib MU, Singh AK, Bhunia AK. Rapid detection and differentiation of Staphylococcus colonies using an optical scattering technology. Anal Bioanal Chem [Internet]. 2018 Sep 1 [cited 2022 Jul 17];410(22):5445–54. Available from: https://pubmed.ncbi.nlm.nih.gov/29796901/

17. Zieliński B, Plichta A, Misztal K, Spurek P, Brzychczy-Włoch M, Ochońska D. Deep learning approach to bacterial colony classification. PLoS One [Internet]. 2017 Sep 1 [cited 2022 Jul 17];12(9):e0184554. Available from: https://journals.plos.org/plosone/article?id=10.1371/journal.pone.0184554

18. Sousa AM, Pereira MO, Lourenço A. MorphoCol: An ontology-based knowledgebase for the characterisation of clinically significant bacterial colony morphologies. J Biomed Inform [Internet]. 2015 Jun 1 [cited 2022 Jul 17];55:55–63. Available from: https://pubmed.ncbi.nlm.nih.gov/25817920/

19. Fratamico PM, Strobaugh TR, Medina MB, Gehring AG. Detection of Escherichia coli 0157:H7 using a surface plasmon resonance biosensor. Biotechnology Techniques 1998 12:7 [Internet]. 1998 [cited 2022 Aug 23];12(7):571–6. Available from: https://link.springer.com/article/10.1023/A:1008872002336

20. Perkins EA, Squirrell DJ. Development of instrumentation to allow the detection of microorganisms using light scattering in combination with surface plasmon resonance. Biosens Bioelectron [Internet]. 2000 Jan [cited 2022 Aug 23];14(10–11):853–9. Available from: https://pubmed.ncbi.nlm.nih.gov/10945460/

21. Turra G, Arrigoni S, Signoroni A. CNN-Based Identification of Hyperspectral Bacterial Signatures for Digital Microbiology. Lecture Notes in Computer Science (including subseries Lecture Notes in Artificial Intelligence and Lecture Notes in Bioinformatics) [Internet]. 2017 [cited 2022 Jul 17];10485 LNCS:500–10. Available from: https://link.springer.com/chapter/10.1007/978-3-319-68548-9_46

22. Andreini P, Bonechi S, Bianchini M, Mecocci A, Scarselli F. A deep learning approach to bacterial colony segmentation. Lecture Notes in Computer Science (including subseries Lecture Notes in Artificial Intelligence and Lecture Notes in Bioinformatics) [Internet]. 2018 [cited 2022 Jul 17];11141 LNCS:522–33. Available from: https://link.springer.com/chapter/10.1007/978-3-030-01424-7_51

23. Geirhos R, Janssen DHJ, Schütt HH, Rauber J, Bethge M, Wichmann FA. Comparing deep neural networks against humans: object recognition when the signal gets weaker. 2017 Jun 21 [cited 2022 Jul 17]; Available from: https://arxiv.org/abs/1706.06969v2

24. Buetti-Dinh A, Galli V, Bellenberg S, Ilie O, Herold M, Christel S, et al. Deep neural networks outperform human expert’s capacity in characterizing bioleaching bacterial biofilm composition. Biotechnology Reports. 2019 Jun 1;22:e00321.

25. Russakovsky O, Deng J, Su H, Krause J, Satheesh S, Ma S, et al. ImageNet Large Scale Visual Recognition Challenge. Int J Comput Vis [Internet]. 2014 Sep 1 [cited 2022 Jul 17];115(3):211–52. Available from: https://arxiv.org/abs/1409.0575v3

26. Deng J, Berg AC, Li K, Fei-Fei L. What Does Classifying More than 10,000 Image Categories Tell Us? In: Proceedings of the 11th European Conference on Computer Vision: Part V. Berlin, Heidelberg: Springer-Verlag; 2010. p. 71–84. (ECCV’10).

27. Russakovsky O, Fei-Fei L. Attribute learning in large-scale datasets. Lecture Notes in Computer Science (including subseries Lecture Notes in Artificial Intelligence and Lecture Notes in Bioinformatics) [Internet]. 2012 [cited 2022 Jul 18];6553 LNCS(PART 1):1–14. Available from: https://collaborate.princeton.edu/en/publications/attribute-learning-in-large-scale-datasets

28. Russakovsky O, Deng J, Huang Z, Berg AC, Fei-Fei L. Detecting avocados to Zucchinis: What have we done, and where are we going? Proceedings of the IEEE International Conference on Computer Vision [Internet]. 2013 [cited 2022 Jul 18];2064–71. Available from: https://collaborate.princeton.edu/en/publications/detecting-avocados-to-zucchinis-what-have-we-done-and-where-are-w

29. Balki I, Amirabadi A, Levman J, Martel AL, Emersic Z, Meden B, et al. Sample-Size Determination Methodologies for Machine Learning in Medical Imaging Research: A Systematic Review. Canadian Association of Radiologists Journal [Internet]. 2019 Nov 1 [cited 2022 Jul 17];70(4):344–53. Available from: https://journals.sagepub.com/doi/10.1016/j.carj.2019.06.002?url_ver=Z39.88-2003&rfr_id=ori%3Arid%3Acrossref.org&rfr_dat=cr_pub++0pubmed

30. Diggle SP, Whiteley M. Microbe profile: Pseudomonas aeruginosa: Opportunistic pathogen and lab rat. Microbiology (United Kingdom) [Internet]. 2020 Oct 10 [cited 2022 Jul 17];166(1):30–3. Available from: https://www.microbiologyresearch.org/content/journal/micro/10.1099/mic.0.000860

31. Moradali MF, Ghods S, Rehm BHA. Pseudomonas aeruginosa Lifestyle: A Paradigm for Adaptation, Survival, and Persistence. Front Cell Infect Microbiol [Internet]. 2017 Feb 15 [cited 2022 Jul 17];7(FEB):39. Available from: /pmc/articles/PMC5310132/

32. Nathwani D, Raman G, Sulham K, Gavaghan M, Menon V. Clinical and economic consequences of hospital-acquired resistant and multidrug-resistant Pseudomonas aeruginosa infections: A systematic review and meta-analysis. Antimicrob Resist Infect Control [Internet]. 2014 Oct 20 [cited 2022 Jul 17];3(1):1–16. Available from: https://aricjournal.biomedcentral.com/articles/10.1186/2047-2994-3-32

33. Elborn JS. Cystic fibrosis. The Lancet. 2016 Nov 19;388(10059):2519–31.

34. Pohl S, Klockgether J, Eckweiler D, Khaledi A, Schniederjans M, Chouvarine P, et al. The extensive set of accessory Pseudomonas aeruginosa genomic components. FEMS Microbiol Lett [Internet]. 2014 Jul 1 [cited 2022 Jul 18];356(2):235–41. Available from: https://academic.oup.com/femsle/article/356/2/235/542446

35. Freschi L, Vincent AT, Jeukens J, Emond-Rheault JG, Kukavica-Ibrulj I, Dupont MJ, et al. The Pseudomonas aeruginosa Pan-Genome Provides New Insights on Its Population Structure, Horizontal Gene Transfer, and Pathogenicity. Genome Biol Evol [Internet]. 2019 Jan 1 [cited 2022 Jul 17];11(1):109–20. Available from: https://pubmed.ncbi.nlm.nih.gov/30496396/

36. Poulsen BE, Yang R, Clatworthy AE, White T, Osmulski SJ, Li L, et al. Defining the core essential genome of Pseudomonas aeruginosa. Proc Natl Acad Sci U S A [Internet]. 2019 May 14 [cited 2022 Jul 17];116(20):10072–80. Available from: /pmc/articles/PMC6525520/

37. Lebreton F, Snesrud E, Hall L, Mills E, Galac M, Stam J, et al. A panel of diverse Pseudomonas aeruginosa clinical isolates for research and development. JAC Antimicrob Resist [Internet]. 2021 Sep 30 [cited 2022 Jul 18];3(4). Available from: https://academic.oup.com/jacamr/article/3/4/dlab179/6458700

38. Kiyaga S, Kyany’a C, Muraya AW, Smith HJ, Mills EG, Kibet C, et al. Genetic Diversity, Distribution, and Genomic Characterization of Antibiotic Resistance and Virulence of Clinical Pseudomonas aeruginosa Strains in Kenya. Front Microbiol. 2022 Mar 14;13:699.

39. Kirisits MJ, Prost L, Starkey M, Parsek MR. Characterization of Colony Morphology Variants Isolated from Pseudomonas aeruginosa Biofilms. Appl Environ Microbiol [Internet]. 2005 Aug [cited 2022 Jul 17];71(8):4809. Available from: /pmc/articles/PMC1183349/

40. Ikeno T, Fukuda K, Ogawa M, Honda M, Tanabe T, Taniguchi H. Small and rough colony pseudomonas aeruginosa with elevated biofilm formation ability isolated in hospitalized patients. Microbiol Immunol [Internet]. 2007 [cited 2022 Jul 17];51(10):929–38. Available from: https://pubmed.ncbi.nlm.nih.gov/17951982/

41. Rakhimova E, Munder A, Wiehlmann L, Bredenbruch F, Tümmler B. Fitness of Isogenic Colony Morphology Variants of Pseudomonas aeruginosa in Murine Airway Infection. PLoS One [Internet]. 2008 Feb 27 [cited 2022 Jul 17];3(2):1685. Available from: /pmc/articles/PMC2246019/

42. Azimi S, Roberts AEL, Peng S, Weitz JS, McNally A, Brown SP, et al. Allelic polymorphism shapes community function in evolving Pseudomonas aeruginosa populations. The ISME Journal 2020 14:8 [Internet]. 2020 Apr 27 [cited 2022 Jul 17];14(8):1929–42. Available from: https://www.nature.com/articles/s41396-020-0652-0

43. Vanderwoude J, Fleming D, Azimi S, Trivedi U, Rumbaugh KP, Diggle SP. The evolution of virulence in Pseudomonas aeruginosa during chronic wound infection. Proceedings of the Royal Society B [Internet]. 2020 Oct 28 [cited 2022 Jul 17];287(1937):20202272. Available from: https://royalsocietypublishing.org/doi/10.1098/rspb.2020.2272

44. Boucher JC, Yu H, Mudd MH, Deretic V. Mucoid Pseudomonas aeruginosa in cystic fibrosis: characterization of muc mutations in clinical isolates and analysis of clearance in a mouse model of respiratory infection. Infect Immun [Internet]. 1997 [cited 2022 Jul 20];65(9):3838–46. Available from: https://pubmed.ncbi.nlm.nih.gov/9284161/

45. Pirnay JP, Bilocq F, Pot B, Cornelis P, Zizi M, van Eldere J, et al. Pseudomonas aeruginosa Population Structure Revisited. PLoS One [Internet]. 2009 Nov 13 [cited 2022 Jul 17];4(11):e7740. Available from: https://journals.plos.org/plosone/article?id=10.1371/journal.pone.0007740

46. Dettman JR, Rodrigue N, Aaron SD, Kassen R. Evolutionary genomics of epidemic and nonepidemic strains of Pseudomonas aeruginosa. Proc Natl Acad Sci U S A [Internet]. 2013 Dec 24 [cited 2022 Jul 17];110(52):21065–70. Available from: www.pnas.org/cgi/doi/10.1073/pnas.1307862110

47. Shrestha SD, Guttman DS, Perron GG. Draft Genome Sequences of 10 Environmental Pseudomonas aeruginosa Strains Isolated from Soils, Sediments, and Waters. Genome Announc [Internet]. 2017 [cited 2022 Jul 17];5(34). Available from: https://journals.asm.org/doi/10.1128/genomeA.00804-17

48. Mikołajczyk A, Grochowski M. Data augmentation for improving deep learning in image classification problem. 2018 International Interdisciplinary PhD Workshop, IIPhDW 2018. 2018 Jun 18;117–22.

49. Wong SC, Gatt A, Stamatescu V, McDonnell MD. Understanding Data Augmentation for Classification: When to Warp? 2016 International Conference on Digital Image Computing: Techniques and Applications, DICTA 2016. 2016 Dec 22;

50. Pratt LY. Discriminability-Based Transfer between Neural Networks. In: Hanson S, Cowan J, Giles C, editors. Advances in Neural Information Processing Systems [Internet]. Morgan-Kaufmann; 1992. Available from: https://proceedings.neurips.cc/paper/1992/file/67e103b0761e60683e83c559be18d40c-Paper.pdf

51. Yu H, Winkler S. Image complexity and spatial information. 2013 5th International Workshop on Quality of Multimedia Experience, QoMEX 2013 - Proceedings. 2013;12–7.

52. Danielsen J, Nordenfelt P. Computer vision-based image analysis of bacteria. Methods in Molecular Biology [Internet]. 2017 [cited 2022 Oct 11];1535:161–72. Available from: https://link.springer.com/protocol/10.1007/978-1-4939-6673-8_10

53. Simonyan K, Zisserman A. Very Deep Convolutional Networks for Large-Scale Image Recognition. 3rd International Conference on Learning Representations, ICLR 2015 - Conference Track Proceedings [Internet]. 2014 Sep 4 [cited 2022 Jul 17]; Available from: https://arxiv.org/abs/1409.1556v6

54. Shaw LP, Wang AD, Dylus D, Meier M, Pogacnik G, Dessimoz C, et al. The phylogenetic range of bacterial and viral pathogens of vertebrates. Mol Ecol [Internet]. 2020 Sep 1 [cited 2022 Jul 17];29(17):3361–79. Available from: https://onlinelibrary.wiley.com/doi/full/10.1111/mec.15463

55. Subedi D, Vijay AK, Kohli GS, Rice SA, Willcox M. Comparative genomics of clinical strains of Pseudomonas aeruginosa strains isolated from different geographic sites. Scientific Reports 2018 8:1 [Internet]. 2018 Oct 23 [cited 2022 Jul 17];8(1):1–14. Available from: https://www.nature.com/articles/s41598-018-34020-7

56. Medina-Rojas M, Stribling W, Snesrud E, Garry BI, Li Y, Gann PM, et al. Comparison of Pseudomonas aeruginosa strains reveals that Exolysin A toxin plays an additive role in virulence. Pathog Dis [Internet]. 2020 Feb 1 [cited 2022 Jul 17];78(1):10. Available from: https://academic.oup.com/femspd/article/78/1/ftaa010/5804881

57. Sharma D, Misba L, Khan AU. Antibiotics versus biofilm: an emerging battleground in microbial communities. Antimicrobial Resistance & Infection Control 2019 8:1 [Internet]. 2019 May 16 [cited 2022 Jul 25];8(1):1–10. Available from: https://aricjournal.biomedcentral.com/articles/10.1186/s13756-019-0533-3

58. Rudin C. Stop explaining black box machine learning models for high stakes decisions and use interpretable models instead. Nature Machine Intelligence 2019 1:5 [Internet]. 2019 May 13 [cited 2022 Jul 20];1(5):206–15. Available from: https://www.nature.com/articles/s42256-019-0048-x

59. Tolstikhin I, Bousquet O, Schölkopf B, Thierbach K, Bazin PL, de Back W, et al. Generative Adversarial Networks. Lecture Notes in Computer Science (including subseries Lecture Notes in Artificial Intelligence and Lecture Notes in Bioinformatics) [Internet]. 2014 Jun 10 [cited 2022 Jul 17];11046 LNCS(NeurIPS):1–9. Available from: https://arxiv.org/abs/1406.2661v1

60. Radford A, Metz L, Chintala S. Unsupervised Representation Learning with Deep Convolutional Generative Adversarial Networks. 4th International Conference on Learning Representations, ICLR 2016 - Conference Track Proceedings [Internet]. 2015 Nov 19 [cited 2022 Jul 17]; Available from: https://arxiv.org/abs/1511.06434v2

61. Cabeen MT, Leiman SA, Losick R. Colony-morphology screening uncovers a role for the Pseudomonas aeruginosa nitrogen-related phosphotransferase system in biofilm formation. Mol Microbiol [Internet]. 2016 Feb 1 [cited 2022 Jul 17];99(3):557. Available from: /pmc/articles/PMC5130288/

62. LeCun Y, Bottou L, Bengio Y, Haffner P. Gradient-based learning applied to document recognition. Proceedings of the IEEE. 1998;86(11):2278–323.

63. Perez L, Wang J. The Effectiveness of Data Augmentation in Image Classification using Deep Learning. 2017 Dec 13 [cited 2023 Jul 11]; Available from: https://arxiv.org/abs/1712.04621v1

64. Alomar K, Aysel HI, Cai X. Data Augmentation in Classification and Segmentation: A Survey and New Strategies. Journal of Imaging 2023, Vol 9, Page 46 [Internet]. 2023 Feb 17 [cited 2023 Jul 11];9(2):46. Available from: https://www.mdpi.com/2313-433X/9/2/46/htm

65. Ioffe S, Szegedy C. Batch Normalization: Accelerating Deep Network Training by Reducing Internal Covariate Shift. CoRR [Internet]. 2015;abs/1502.0. Available from: http://arxiv.org/abs/1502.03167

66. Shorten C, Khoshgoftaar TM. A survey on Image Data Augmentation for Deep Learning. J Big Data [Internet]. 2019 Dec 1 [cited 2023 Jul 11];6(1):1–48. Available from: https://journalofbigdata.springeropen.com/articles/10.1186/s40537-019-0197-0

67. Cubuk ED, Zoph B, Mané D, Vasudevan V, Le Q V. AutoAugment: Learning Augmentation Policies from Data. CoRR [Internet]. 2018;abs/1805.0. Available from: http://arxiv.org/abs/1805.09501

68. Nair V, Hinton GE. Rectified Linear Units Improve Restricted Boltzmann Machines. In: International Conference on Machine Learning. 2010.

69. Yosinski J, Clune J, Bengio Y, Lipson H. How transferable are features in deep neural networks? CoRR [Internet]. 2014;abs/1411.1. Available from: http://arxiv.org/abs/1411.1792

70. He K, Zhang X, Ren S, Sun J. Deep Residual Learning for Image Recognition. CoRR [Internet]. 2015;abs/1512.0. Available from: http://arxiv.org/abs/1512.03385

71. Alhammad S, Zhao K, Jennings A, Hobson P, Smith DF, Baker B, et al. Efficient DNN-Based Classification of Whole Slide Gram Stain Images for Microbiology. In: 2021 Digital Image Computing: Techniques and Applications (DICTA). 2021. p. 1–8.

72. Shwetha V, Prasad K, Mukhopadhyay C, Banerjee B, Chakrabarti A. Automatic Detection of Bacilli Bacteria from Ziehl-Neelsen Sputum Smear Images. In: 2021 2nd International Conference on Communication, Computing and Industry 40 (C2I4). 2021. p. 1–5.

73. Sandler M, Howard AG, Zhu M, Zhmoginov A, Chen LC. Inverted Residuals and Linear Bottlenecks: Mobile Networks for Classification, Detection and Segmentation. CoRR [Internet]. 2018;abs/1801.0. Available from: http://arxiv.org/abs/1801.04381

74. Chollet F. Xception: Deep Learning with Depthwise Separable Convolutions. CoRR [Internet]. 2016;abs/1610.0. Available from: http://arxiv.org/abs/1610.02357

75. Chen Y, Yu G, Long Y, Teng J, You X, Liao BQ, et al. Application of radial basis function artificial neural network to quantify interfacial energies related to membrane fouling in a membrane bioreactor. Bioresour Technol [Internet]. 2019;293:122103. Available from: https://www.sciencedirect.com/science/article/pii/S0960852419313331

76. Rahmayuna N, Rahardwika D, Sari A, Setiadi DRIM, Rachmawanto E. Pathogenic Bacteria Genus Classification using Support Vector Machine. In 2018. p. 23–7.

